# Multiple NTS Neuron Populations Synergistically Suppress Physiologic Food Intake but are Dispensable for the Response to VSG

**DOI:** 10.1101/2022.12.23.521804

**Authors:** Weiwei Qui, Chelsea R. Hutch, Yi Wang, Jennifer Wloszek, Rachel A. Rucker, Martin G. Myers, Darleen Sandoval

## Abstract

Several discrete groups of feeding-regulated neurons in the *nucleus tractus solitarius* (NTS) suppress food intake, including aversion-promoting neurons that express *Cck* (NTS^Cck^ cells) and distinct *Lepr*- and *Calcr*-expressing neurons (NTS^Lepr^ and NTS^Calcr^ cells, respectively) that suppress food intake without promoting aversion. To test synergies among these cell groups we manipulated multiple NTS cell populations simultaneously. We found that activating multiple sets of NTS neurons (e.g., NTS^Lepr^ plus NTS^Calcr^ (NTS^LC^), or NTS^LC^ plus NTS^Cck^ (NTS^LCK^)) suppressed feeding more robustly than activating single populations. While activating groups of cells that include NTS^Cck^ neurons promoted conditioned taste avoidance (CTA), NTS^LC^ activation produced no CTA despite abrogating feeding. Thus, the ability to promote CTA formation represents a dominant effect, but activating multiple non-aversive populations additively suppresses food intake without provoking aversion. Although silencing multiple NTS neuron groups augmented food intake and body weight more dramatically than silencing single populations, feeding activated many non-NTS^LCK^ neurons and silencing NTS^LCK^ neurons failed to blunt the weight loss response to vertical sleeve gastrectomy (VSG). Hence, while each of these NTS neuron populations plays crucial and additive roles in the control of energy balance, as-yet undefined cell types must make additional contributions to the control of feeding and the response to VSG.

## Introduction

The ongoing obesity pandemic represents an enormous challenge to human health and longevity (1). Identifying new therapeutic targets to better combat obesity will require understanding the physiologic systems that modulate feeding and contribute to body weight maintenance. While hypothalamic circuits contribute to the control of feeding and play important roles in maintaining long-term energy balance, many hypothalamic circuits act via the brainstem to suppress food intake (2, 3). Furthermore, the brainstem receives feeding-related and other information from the gastrointestinal (GI) tract, controls the rhythmic pattern generators that mediate feeding, and can overcome hypothalamically-driven hyperphagia (2, 4).

Many of the most important food intake-controlling brainstem systems lie in the dorsal vagal complex (DVC), which includes the area postrema (AP), the *nucleus tractus solitarius* (NTS), and the dorsal motor nucleus of the vagus (DMV) (2, 3). The AP lies outside of the blood-brain barrier and receives circulating signals relevant to feeding, including from several anorexigenic peptides. The NTS, which lies adjacent to and receives input from the AP, also receives input from GI-innervating vagal sensory neurons. Both the AP and NTS project ventrally to the DMV to control GI motility and other physiological functions, in addition to innervating more rostral brain regions to modulate food intake.

Many of these brainstem circuits express receptors for peptides that suppress food intake, including the glucagon-like peptide-1 receptor (GLP1R); the calcitonin receptor (CALCR) and the related amylin receptor (AmyR-a complex of CALCR and a receptor activity modifying protein (RAMP)); and the leptin receptor (LepRb) (2, 5-7). Many of these DVC cell types may contribute to the suppression of feeding by a variety of gut peptide mimetics currently under development or in use for weight loss therapy (2). Furthermore, recent work from others and us has demonstrated that the AP and NTS neuron types that suppress food intake while mediating aversive responses associated with gut malaise are distinct from neuron types that promote non-aversive satiation (6-11). Cell types that suppress food intake without causing aversion include NTS neurons that express *Lepr* (NTS^Lepr^ cells) or *Calcr* (NTS^Calcr^ cells); in contrast, *Cck*-expressing NTS neurons (NTS^Cck^ cells) promote aversion (4, 6, 11).

Bariatric surgeries, including vertical sleeve gastrectomy (VSG), decrease meal size and overall food intake to promote dramatic weight loss in many patients (12), as well as in animal models of VSG (13). The irreversibility and side effects of the procedure, together with the inability to perform enough surgeries to treat the number of patients in need, prevents the use of this intervention at a population scale, however (14). Hence, we must understand the mechanisms by which VSG mediates its effects to help identify therapeutic targets with which to medically mimic its actions.

To determine the potential additivity of food intake-regulating NTS circuits and the potential roles for these circuits in the response to VSG, we simultaneously manipulated multiple NTS cell types. We found that activating multiple non-aversive cell types fails to promote aversive effects despite abrogating food intake. Furthermore, silencing multiple NTS cell populations augmented weight gain in high fat diet (HFD)-fed mice but did not alter weight loss in response to VSG. Thus, while each of these NTS neuron populations plays crucial and independent roles in the control of energy balance, as-yet undefined cell types must make additional contributions to the control of feeding, as well as mediating the response to VSG.

## Results

### Additive effects on food intake suppression by NTS^Lepr^ and NTS^Calcr^ neurons

We have previously shown that NTS^Lepr^ and NTS^Calcr^ cells represent distinct populations of NTS neurons (5, 6), and that the activation of either population mediates the non-aversive suppression of food intake (6, 11). To determine the relative ability of each of these cell types to mediate the suppression of food intake and to determine whether they might additively suppress feeding, we injected an adeno-associated virus (AAV) to cre-dependently express the activating (hM3Dq) designer receptor exclusively activated by designer drugs (DREADD; AAV^hM3Dq^ (15, 16)) into the NTS of *Lepr*^*Cre*^, *Calcr*^*Cre*^, or dual *Lepr*^*Cre*^*;Calcr*^*Cre*^ mice (producing Lepr^Dq^, Calcr^Dq^, and LC^Dq^ mice, respectively). We then examined the effects of CNO-mediated hM3Dq activation on food intake and related parameters in these mice (Figure 1). As expected, CNO treatment promoted FOS-immunoreactivity (-IR) in the NTS of Lepr^Dq^, Calcr^Dq^ and LC^Dq^ mice, with the LC^Dq^ mice displaying increased FOS-IR compared to animals in which we had transduced only one population (Figure 1A).

**Figure 1:**
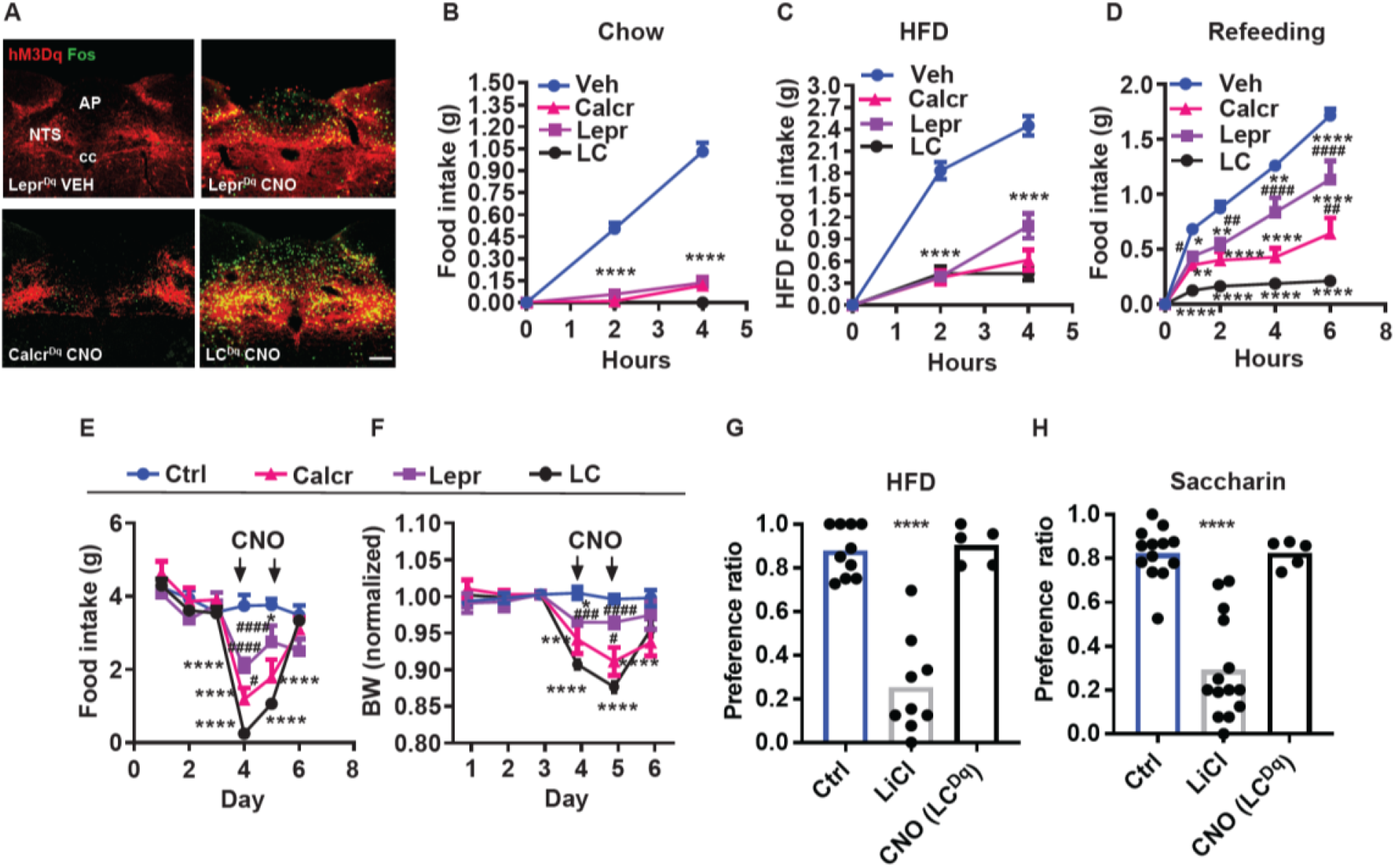
Additive suppression of food intake with combined activation of NTS^Calcr^ and NTS^Lepr^ neurons. (**A**) Representative images showing mCherry-IR (hM3Dq, red) and FOS-IR (green) in the NTS of Lepr^Dq^, Calcr^Dq^ and LC^Dq^ mice following treatment with saline (Lepr^Dq^ VEH) or CNO (Lepr^Dq^ CNO, Calcr^Dq^ CNO and LC^Dq^ CNO IP, 1 mg/kg) for 2 hours before perfusion. NTS: nucleus of the solitary tract, AP: area postrema, cc: central canal. All images were taken at the same magnification; scale bar equals 150 μm. (**B-D**) Food intake in chow-fed Lepr^Dq^ (Lepr), Calcr^Dq^ (Calcr) and LC^Dq^ (LC) mice over the first 4 hours of the dark phase following vehicle (Veh) or CNO injection (IP, 1 mg/kg) when provided with chow (**B**, n=25 for Veh group; n=20 for both Lepr and Calcr groups; n=8 for LC group) or HFD (**C**, n=21 for Veh group; n=6 for Lepr group; n=7 for Calcr group; n=8 for LC group). (**D**) Food intake for the same groups of mice over the first 6 hours of refeeding during the light cycle following an overnight fast (**D**, n=23 for Veh group; n=8 for Lepr group; n=7 for Calcr group; n=7 for LC group) following with CNO (IP, 1 mg/kg) or vehicle (Veh). (**E-F**) Control (Ctrl, n=5) or Lepr^Dq^ (Lepr, n=5), Calcr^Dq^ (Calcr, n=5) and LC^Dq^ (LC, n=8) mice were treated with vehicle for three baseline days, followed by two days of twice daily treatment with CNO (IP 1 mg/kg), followed by two additional days of Veh treatment. Daily food intake (**E**) and body weight (shown as change from baseline) (**F**). Vehicle and CNO treatment are denoted on the graphs. (**G, H**) CTA assays: Mice were treated with vehicle (Veh, n=10), LiCl (IP, 126 mg/kg, n=9), or CNO (IP, 1 mg/kg, n=5) during exposure to a novel tastant (HFD) (**G**) or saccharin (**H**) paired with vehicle (Veh, n=13), LiCl (IP, 126 mg/kg, n=14), or CNO (IP, 1 mg/kg, n=5). Shown is mean +/-SEM; Two-way ANOVA, sidak’s multiple comparisons test was used; *p<0.05, **p<0.01, ***p<0.001, ****p<0.0001 vs vehicle or Ctrl; ^#^p<0.05, ^##^p<0.01, ^###^p<0.001, ^####^p<0.0001 vs LC.

As we showed previously, CNO almost completely abrogated food intake at the onset of the dark cycle in chow-fed Lepr^Dq^ and Calcr^Dq^ mice; LC^Dq^ mice responded similarly (Figure 1B). The same was true for mice fed high-fat diet (HFD) at the onset at the dark cycle, although the activation of NTS^Lepr^ neurons tended to suppress feeding less than NTS^Calcr^ and NTS^LC^ neurons at the 4-hour time point in this assay (Figure 1C). Following an overnight fast, the activation of NTS^Lepr^ neurons significantly decreased refeeding over 6 hours; the activation of NTS^Calcr^ cells alone decreased refeeding further than NTS^Lepr^ neurons, and the activation of NTS^LC^ neurons decreased refeeding even further-by approximately 90% (Figure 1D).

Similarly, activating NTS^Lepr^ neurons decreased feeding and body weight over a two-day treatment, activating NTS^Calcr^ neurons suppressed these parameters more effectively, and activating both cell types in LC^Dq^ mice essentially abolished feeding for the first 24 hours and continued to reduce food intake compared to the individual cell types during the second 24 hours of the experiment (Figure 1 E, F). While we had intended to continue treatment for another day, we discontinued CNO treatment after 48 hours because of animal welfare concerns due to the amount of weight loss exhibited by the LC^Dq^ mice. Hence, NTS^Calcr^ neurons suppress feeding more robustly than NTS^Lepr^ neurons, and NTS^Calcr^ and NTS^Lepr^ neurons suppress feeding additively.

To determine whether the dramatic decrease in feeding promoted by NTS^LC^ neuron activation provoked aversive responses, we examined whether CNO treatment in LC^Dq^ mice would provoke a CTA. We found that, although the gut malaise associated with LiCl injection promoted a robust CTA to either the novel exposure to HFD or saccharine in drinking water, NTS^LC^ activation failed to provoke such a response (Figure 1G, H). Thus, the coordinated activation of two non-aversive cell types (NTS^Lepr^ and NTS^Calcr^ (6, 11, 17)) fails to promote a CTA despite abolishing food intake.

We also expressed the inhibitory (hM4Di) DREADD in NTS^LC^ cells to examine the effect of acutely inhibiting these cells on short-term food intake (Supplemental Figure 1). While CNO failed to significantly increase food intake during the first four hours of the dark cycle, stimulating hM4Di in NTS^LC^ cells increased food intake during refeeding with chow or HFD following an overnight fast. Thus, NTS^LC^ neurons contribute to the restraint of food intake during refeeding.

We previously showed that the long-term tetanus toxin (TetTox)-mediated silencing of NTS^Calcr^ neurons promotes increased food intake and body weight in animals exposed to HFD (6). To determine whether silencing NTS^Lepr^ cells might mediate similar effects (thereby implicating this additional NTS cell type in the control of long-term energy balance) we injected an AAV to cre-dependently express TetTox into the NTS of *Lepr*^*Cre*^ mice (AAV^TetTox^ (18); Lepr^TetTox^ mice) and examined their food intake and body weight during 6 weeks of chow feeding followed by an additional 6 weeks of HFD feeding (Figure 2A-D). While we detected no significant increase in food intake or body weight during chow feeding for Lepr^TetTox^ mice, these animals displayed a modest increase in food intake on HFD and gained more body weight over 6 weeks than control animals. Thus, NTS^Lepr^ neurons (like NTS^Calcr^ cells) contribute to the long-term control of energy balance.

**Figure 2:**
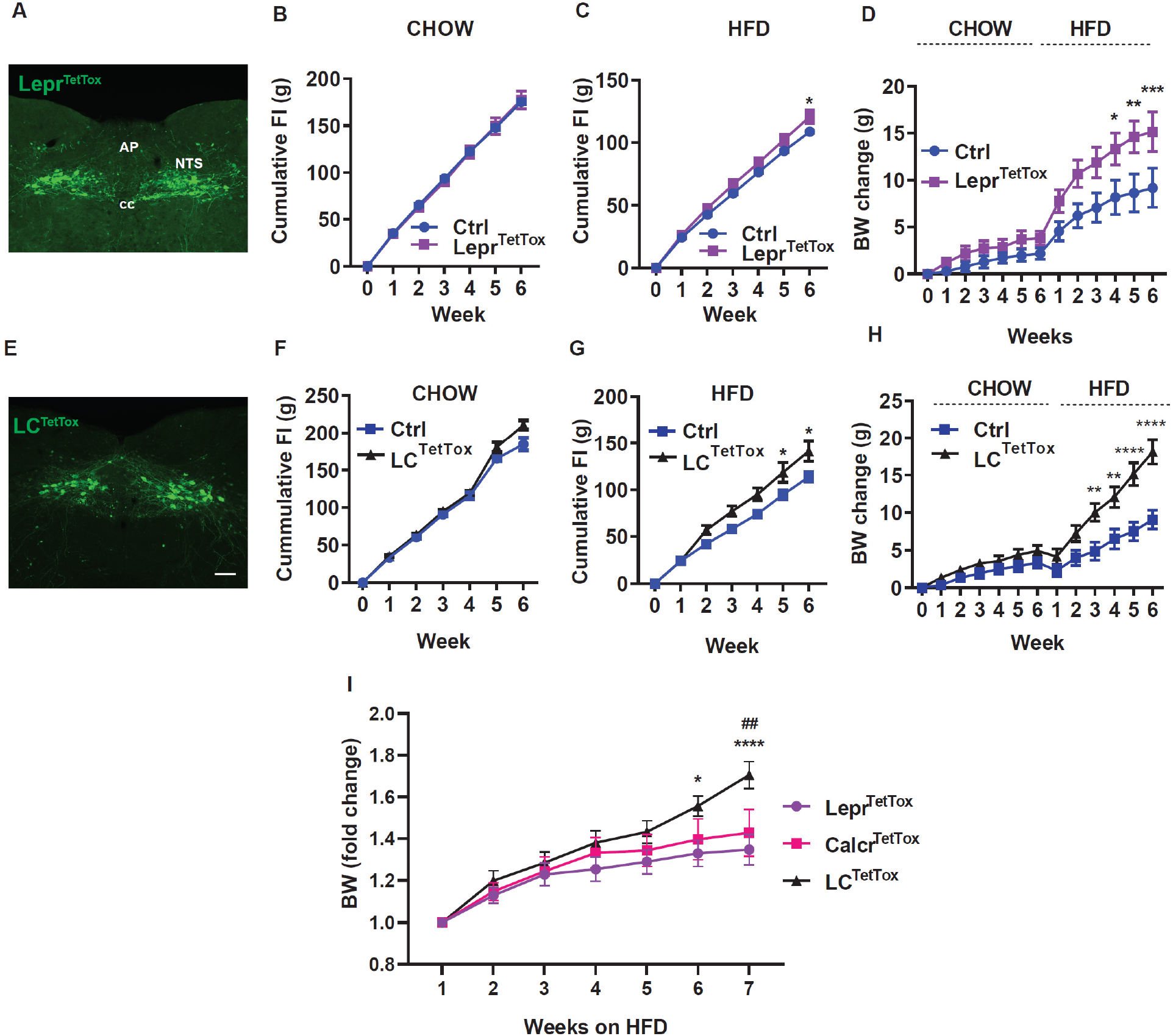
Silencing Lepr^NTS^ neurons or LC^NTS^ neurons exacerbates diet-induced obesity (DIO). Representative images showing GFP-IR (green) from in Lepr^TetTox^ (**A**) and LC^TetTox^ (**E**) mice. All images were taken at the same magnification; scale bar equals 150 μm. NTS: nucleus of the solitary tract, AP: area postrema, cc: central canal. **B-D, F-H** show cumulative food intake during chow (**B, F**) and HFD feeding (**C, G**) and body weight (change from baseline) (**D, H**) for control (Ctrl), Lepr^TetTox^ (**B-D**) and LC^TetTox^ (**F-H**) mice following VSG, during which time they were fed with chow for 5-6 weeks and HFD for an additional 6 weeks. n=7 each group for **B** and **C**, n=8 each group for **D** for Ctrl and Lepr^TetTox^ group; n=7 for Ctrl and n=8 for LC^TetTox^ group in **F** and **G**, n=11 for Ctrl and n=12 for LC^TetTox^ group in **H**. (**I**) Shows weight gain from the onset of HFD feeding for Lepr^TetTox^ (from **D**) and LC^TetTox^ mice (from **H**) in comparison to previously-published Calcr^TetTox^ mice (6). Shown are mean +/-SEM. Two-way ANOVA, sidak’s multiple comparisons test was used; *p<0.05, **p<0.01, ***p<0.001, ****p<0.0001 vs Ctrl, except in **I**, where these refer to the comparison between Lepr^TetTox^ and LC^TetTox^; ^##^p<0.01 between Calcr^TetTox^ and LC^TetTox^.

To determine whether silencing NTS^Calcr^ neurons might exacerbate the increased food intake and body weight displayed by Lepr^TetTox^ mice, we used AAV^TetTox^ to cre-dependently express TetTox in the NTS of *Lepr*^*Cre*^*;Calcr*^*Cre*^ animals (LC^TetTox^ mice) (Figure 2E-H). While we detected no significant increase in food intake or body weight during 6 weeks of chow feeding in LC^TetTox^ mice compared to controls, LC^TetTox^ mice ate significantly more food and gained (∼10 grams) more body weight than controls when exposed to HFD for 6 weeks.

Continuous analysis of feeding demonstrated increased food intake by LC^TetTox^ mice in the dark (active) cycle, consistent with decreased satiety signaling (Supplemental Figure 2). While comparing results among different strains of mice studied at different times requires cautious interpretation, these findings suggest that the silencing of multiple NTS neuron populations (as in LC^TetTox^ mice) increases feeding and weight gain relative to silencing single populations (NTS^Lepr^ in the current study, as well as previously-studied NTS^Calcr^ cells previously (6)(Figure 2I)).

Given the dramatic effects on food intake and body weight observed upon silencing NTS^LC^ cells, we decided to test the requirement for the function of these cells in the weight loss response to VSG. We thus generated HFD-fed control and LC^TetTox^ animals as in Figure 2 and subjected them to VSG (Supplemental Figure 2). While, as expected, HFD-fed LC^TetTox^ mice weighed more than HFD-fed control animals at the time of surgery, VSG also promoted more weight loss in these animals than in controls (both in terms of total weight lost and percent body weight lost) and similarly improved measures of glycemic control. Thus, the action of NTS^LC^ neurons is not required for VSG-mediated weight loss.

### Additive roles for aversive and non-aversive NTS cell types for the control of energy balance

While NTS^Lepr^ and NTS^Calcr^ cells both mediate the non-aversive suppression of food intake, activation of NTS^Cck^ neurons promotes aversive (CTA) responses, as well as decreasing feeding (6, 8, 11, 19). Thus, to understand potential roles for aversive NTS^Cck^ cells in the control of energy balance, we injected cre-dependent AAV^TetTox^ into the NTS of *Cck*^*Cre*^ mice (Cck^TetTox^ mice; Figure 3A) and examined their food intake and body weight during 6 weeks of chow feeding followed by an additional 6 weeks of HFD feeding (Figure 3B-D). While we detected no significant increase in food intake or body weight during chow feeding for Cck^TetTox^ mice, these animals gained more body weight over 6 weeks than control animals during HFD feeding. Thus, while it is possible that the inaccuracies inherent to food intake measurement prevented us from detecting an alteration in food intake in the Cck^TetTox^ animals, silencing NTS^Cck^ neurons might impact body weight by other means, including by altering calorie absorption or energy expenditure. In any case, NTS^Cck^ neurons, like NTS^Calcr^ and NTS^Lepr^ cells, contribute to the long-term control of energy balance.

**Figure 3:**
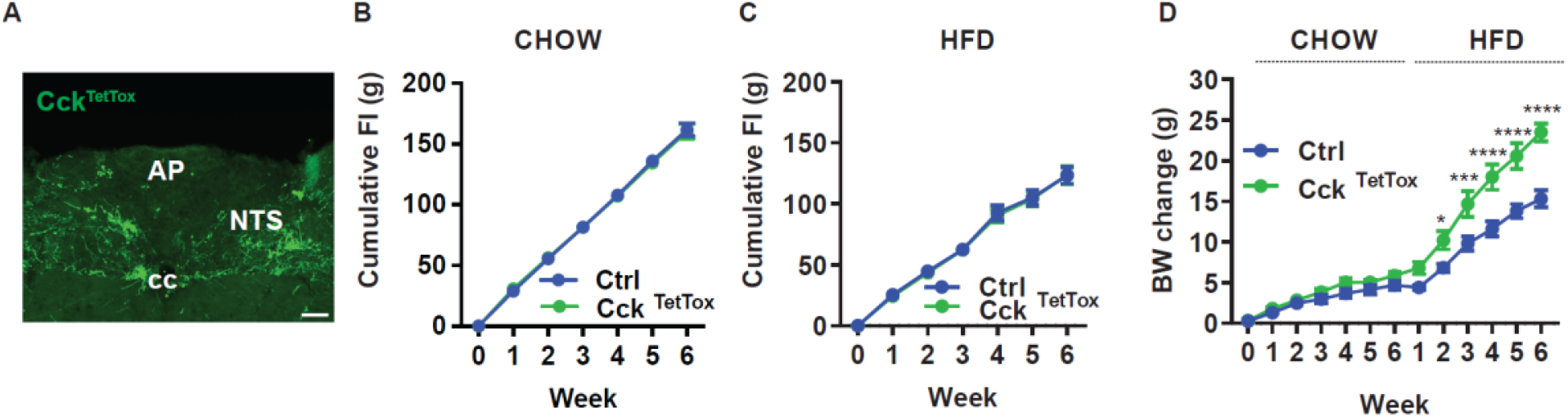
Silencing Cck^NTS^ neurons exacerbates DIO. (**A**) Representative images showing GFP-IR (green) in the NTS of Cck^TetTox^ mice. Images were taken at the same magnification; scale bar equals 150 μm. NTS: nucleus of the solitary tract, AP: area postrema, cc: central canal. (**B-D**) Cumulative food intake during chow (**B**) and HFD (**C**) feeding and body weight (**D**, change from baseline) is shown for control (Ctrl) and Cck^TetTox^ mice following surgery, during which time they were fed with chow for 6 weeks and the HFD for an additional 6 weeks. n=5 each group for **B**, n=6 for Ctrl and n=7 for Cck^Tettox^ group in **C**, n=11 for Ctrl group and n=10 for Cck^TetTox^ group in **D**. Shown is mean +/-SEM. Two-way ANOVA, sidak’s multiple comparisons test was used, *p<0.05, ****p<0.0001 vs Ctrl.

To determine whether aversive NTS^Cck^ neurons might play a role in the weight loss response to VSG, we decided to test whether the function of these cells for the weight loss response to VSG. We thus generated HFD-fed control and Cck^TetTox^ animals as in Figure 3 and subjected them to VSG (Supplemental Figure 3). While, as expected, HFD-fed Cck^TetTox^ mice weighed more than HFD-fed control animals at the time of surgery, VSG promoted similar weight loss in these animals as in controls.

To understand the potential additivity of effects on food intake for aversive and non-aversive NTS cell types, we chose to examine the effects of manipulating NTS^Cck^ neurons (6, 8, 19) in combination with NTS^LC^ cells (collectively, NTS^LCK^ cells) (Figures 4-5). We began by injecting cre-dependent AAV^hM3Dq^ (15, 16) into the NTS of *Lepr*^*Cre*^*;Calcr*^*Cre*^*;Cck*^*Cre*^ mice (producing LCK^Dq^ mice) (Figure 4). We then examined the effects of CNO-mediated activation of NTS^LCK^ cells on food intake and related parameters in these animals.

**Figure 4:**
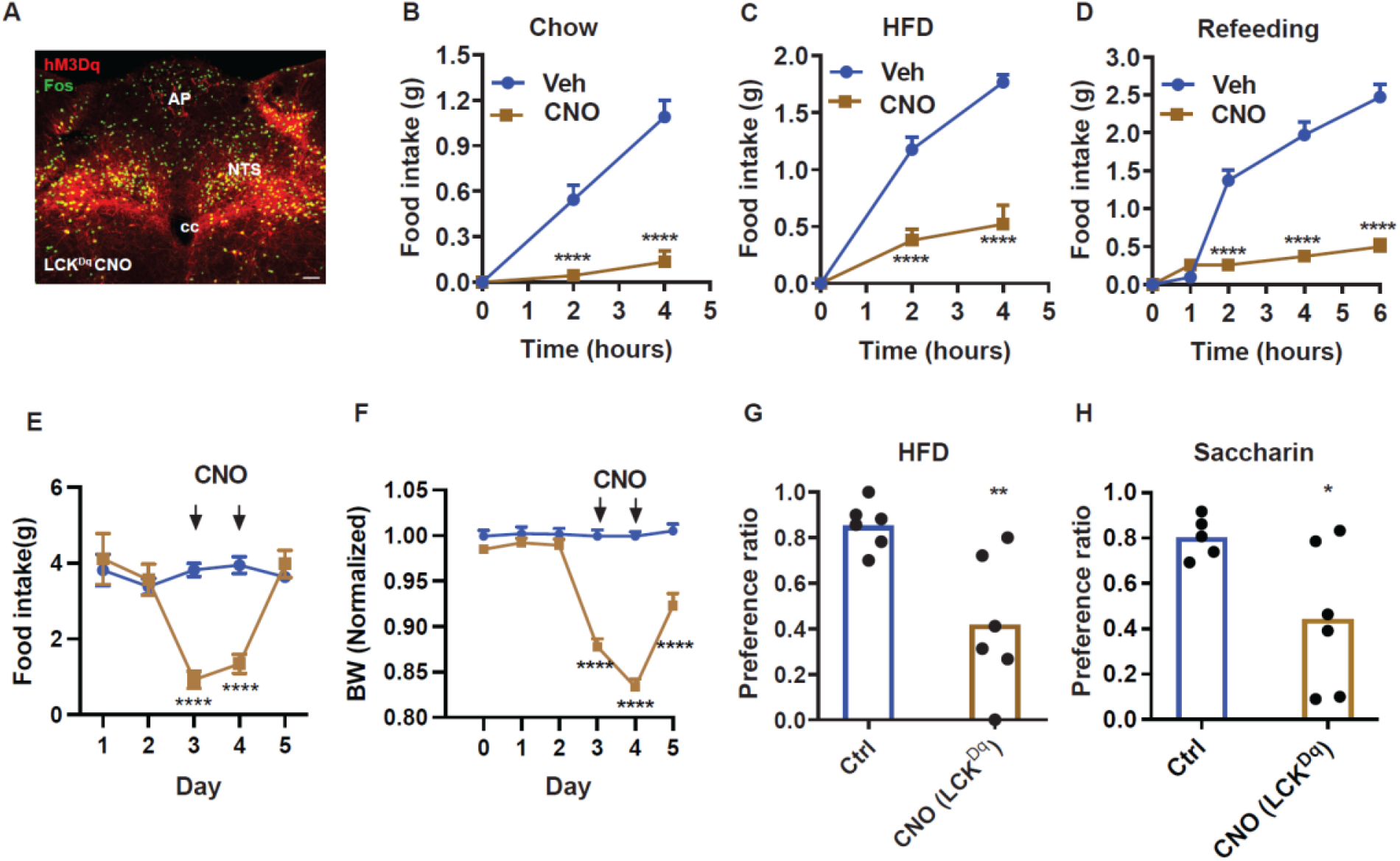
LCK^NTS^ neuron activation decreases food intake and promotes a CTA. (**A**) Representative images showing mCherry-IR (hM3Dq, red) and FOS-IR (green) in the NTS of LCK^Dq^ mice following treatment with CNO (IP, 1 mg/kg) 2 hours before perfusion. Scale bar equals 150 μm. NTS: nucleus of the solitary tract, AP: area postrema, cc: central canal. (**B-D**) Food intake in LCK^Dq^ mice over the first 4 hours of the dark phase during expose to chow (**B**) or HFD (**C**) (n=9 per group), and during the first 6 hours of refeeding in the light cycle following an overnight fast (**D**, n=7 per group) during treatment with CNO (IP, 1 mg/kg) or vehicle. (**E-F**) Control (Ctrl, n=7) or LCK^Dq^ mice (n=7) mice were treated vehicle (Veh) for one days, followed by two days with CNO (1 mg/kg, IP, twice per day) and daily food intake (**E**) and body weight (change from baseline) (**F**) were determined. Veh and CNO treatment are denoted on the graphs. (**G, H**) Mice were treated with Veh, LiCl, or CNO (IP, 1 mg/kg) during exposure to a novel tastant (HFD (**G**) or saccharin (**H**)); n=6 per group. Shown is mean +/-SEM; Two-way ANOVA, sidak’s multiple comparisons test was used in all figures, except panels G and H where unpaired T test was used; *p<0.05, **p<0.01, ***p<0.001, ****p<0.0001 vs Veh or Ctrl.

As expected, CNO treatment promoted FOS-IR in the NTS of LCK^Dq^ mice (Figure 4A). As for LC^Dq^ mice (Figure 1), CNO almost completely abrogated food intake at the onset of the dark cycle in chow-fed and HFD-fed LCK^Dq^ mice, and strongly suppressed refeeding following an overnight fast (Figure 4B-D). We also assessed the effect of activating NTS^LCK^ neurons over a two-day treatment. CNO treatment dramatically decreased food intake in LCK^Dq^ mice and resulted in even larger decreases in body weight that in LC^Dq^ mice (Figures 1E-F, 4E-F), confirming the effectiveness of the coordinated action of NTS^LCK^ neurons for the control of food intake and body weight. Because food intake during CNO treatment was similarly low for LC^Dq^ and LCK^Dq^ mice, while the LCK^Dq^ mice lost more body weight, it is possible that the activation of NTS^Cck^ neurons alters energy balance by altering energy absorption or expenditure, as well as by decreasing feeding. We also found that, unlike NTS^LC^ cells, activating NTS^LCK^ cells provided a strong CTA to HFD or saccharine in water, consistent with the aversive nature of NTS^Cck^ neurons and the dominant nature of aversive over non-aversive hindbrain signals (Figure 4G-H).

To determine whether silencing NTS^Cck^ neurons might exacerbate the increased food intake and body weight displayed by LC^TetTox^ mice (and vice-versa), we injected cre-dependent AAV^TetTox^ into the NTS of *Lepr*^*Cre*^*;Calcr*^*Cre*^*;Cck*^*Cre*^ animals (LCK^TetTox^ mice) (Figure 5A). While we had not detected increased food intake on chow diet in Lepr^TetTox^, Calcr^TetTox^, Cck^TetTox^, or LC^TetTox^ animals (Figures 2-3), LCK^TetTox^ mice displayed a significant increase in food intake on chow, as well as on HFD (Figure 5B-C). Although LCK^TetTox^ mice also tended to gain more weight on chow diet, the effect was not statistically significant; these animals became significantly heavier than controls after the first week on HFD, however, and continued to gain weight relative to controls with each additional week (Figure 5D).

**Figure 5:**
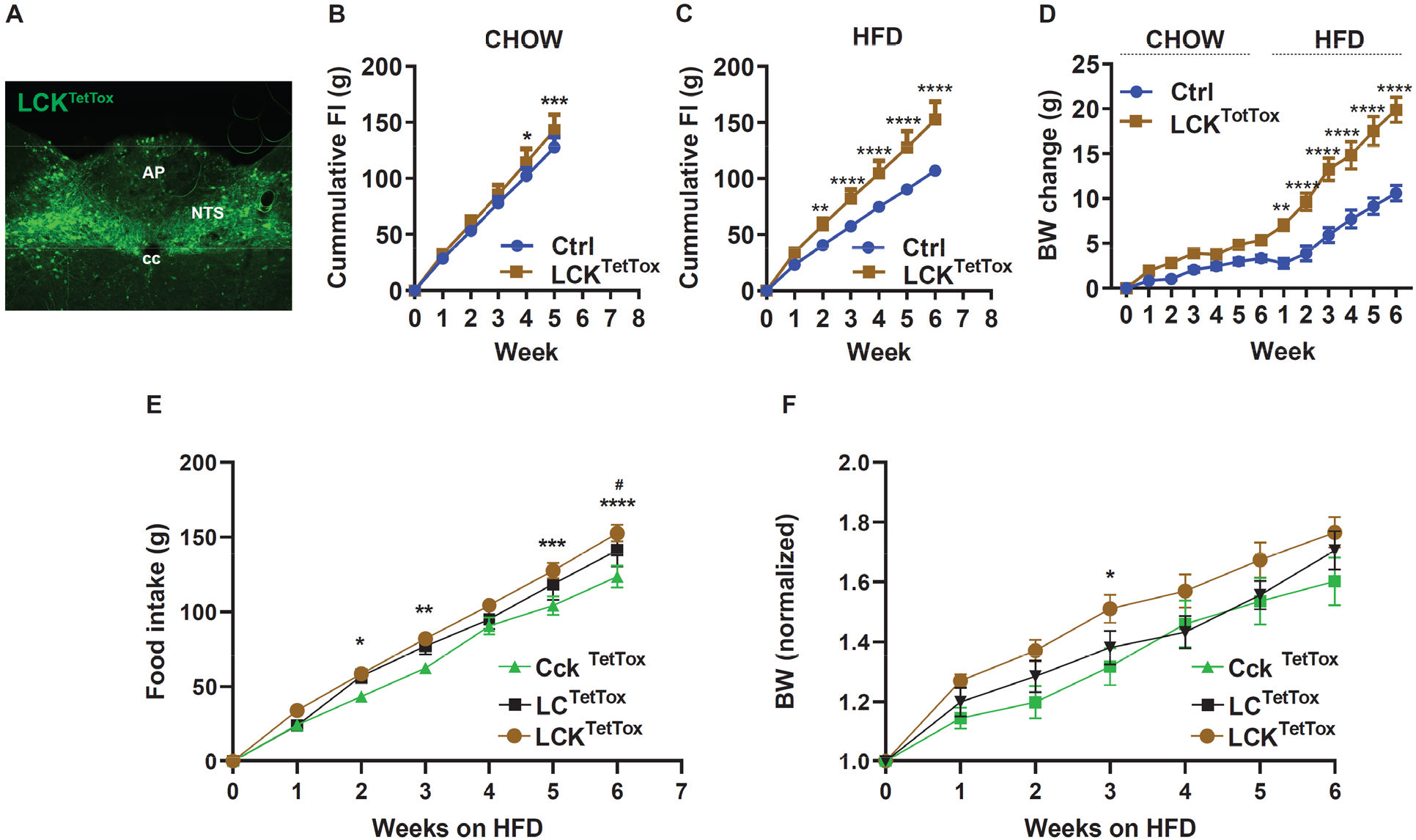
Silencing of LCK^NTS^ neurons increases food intake and body weight. **(A)** Representative image showing GFP-IR (green) in the NTS of LCK^TetTox^ mice. Scale bar equals 150 μm. NTS: the nucleus of the solitary tract, AP: area postrema, cc: central canal. (**B-D**) Cumulative food intake in response to chow (**B**, n=8 per group) and HFD (**C**, n=5 for Ctrl group and n=8 for LCK^TetTox^ group) and body weight (**D**, change from baseline; n=6 for Ctrl group and n=8 for LCK^TetTox^ group) is shown for control (Ctrl) and LCK^TetTox^ mice following surgery, during which time they were fed chow for 6 weeks and HFD for an additional 6 weeks. (**E-F**) Comparisons of cumulative HFD food intake (**E**) and body weight (normalized to baseline) for Cck^TetTox^ (from Figure 3C), LC^TetTox^ (from Figure 2G) and LCK^TetTox^ (from panel **C**) mice. Shown is mean +/-SEM; Two-way ANOVA, sidak’s multiple comparisons test was used; *p<0.05, **p<0.01, ***p<0.001, ****p<0.0001 vs Ctrl except **E** and **F**, for which these indicate differences between Cck^TetTox^ and LCK^TetTox^; ^#^p<0.05 for Cck^TetTox^ vs LC^TetTox^.

While one must exercise caution when comparing results among different strains of mice, we observed increased HFD food intake for LCK^TetTox^ animals compared to Cck^TetTox^ and LC^TetTox^ mice (Figure 5E). Similarly, LCK^TetTox^ mice tended to gain more weight on HFD compared to LC^TetTox^ and Cck^TetTox^ animals (Figure 5F). These findings suggest that simultaneously silencing multiple food intake-controlling NTS neuron populations additively increases food intake and body weight, irrespective of whether the neurons mediate aversive or non-aversive responses.

### NTS^LCK^ neuron signaling is not required for the response to VSG

Given the dramatic effects on food intake and body weight observed upon silencing NTS^LCK^ cells, we decided to test the requirement for the combined function of these cells for the weight loss response to VSG. We thus generated HFD-fed control and LCK^TetTox^ animals as in Figure 5 and subjected them to sham or VSG surgery (Figure 6). While, as expected, HFD-fed LCK^TetTox^ mice weighed more than HFD-fed control animals at the time of surgery, VSG promoted more body and fat mass loss in these animals than in controls, both in terms of total weight lost and percent body weight lost (Figure 6A-B). Indeed, 5-6 weeks after VSG, fat mass for LCK^TetTox^ animals became similar to that of control VSG animals (Figure 6C-E).

**Figure 6:**
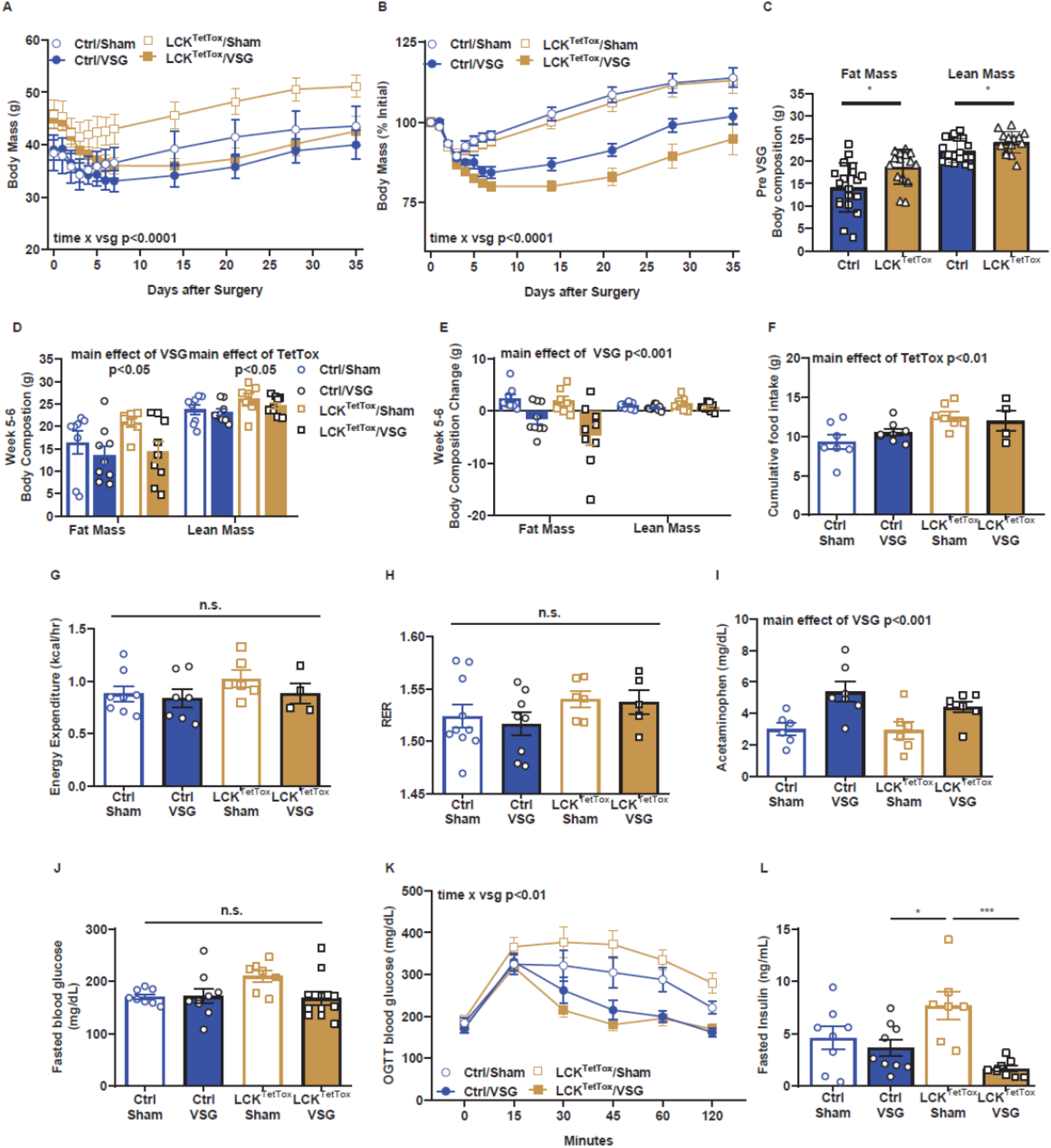
Appropriate response to VSG for LCK^TetTox^ mice. (**A-E**) Shown is body weight (absolute (**A**) and %initial (**B**)) and body composition before (**C**) and 6 weeks after (**D**) VSG, along with change in body composition from baseline (**E**) for body mass in control and LCK^TetTox^ groups after VSG or sham surgery. (**F-H**) A subset of mice was placed in metabolic chambers for 7 days (3 days acclimation, 4 days data analysis) for the determination of food intake (**F**, cumulative over the 4-day data collection period) energy expenditure (**G**), and RER (**H**). (**I**) Gastric emptying rate assessed by acetaminophen appearance in the plasma following gavage. (**J-L**) Fasting blood glucose (**J**), oral glucose tolerance (2 g/kg) (**K**), and fasting insulin are shown for all groups. Data were analyzed using RM-ANOVA or Student’s T-test and are reported as mean +/-SEM. Males mice ctrl/sham (n=6-8), ctrl/VSG (n=7-9), LCK^TetTox^/sham (n=6-7), LCK^TetTox^/VSG (n=4-9). *p<0.05, **p<0.01, ***p<0.001, ****p<0.0001 for the indicated comparisons.

Food intake was decreased by VSG in LCK^TetTox^ animals, although it remained higher than for control mice that had received VSG (Figure 6F)-presumably reflecting the increased food intake of these mice at baseline. We also studied these animals in metabolic cages, which revealed no significant change in metabolic rate or respiratory exchange ratio among groups (Figure 6G-H).

Because the hindbrain receives input from the gut and regulates gastric emptying, we assessed the gastric emptying rate by administering acetaminophen via gavage and measuring its appearance in the plasma. We found that VSG increased gastric emptying rate similarly in control and LCK^TetTox^ mice compared to sham controls (Figure 6I). Glycemic excursions during an oral glucose tolerance test (OGTT) were decreased (improved) and fasting insulin was decreased in LCK^TetTox^ VSG mice in a manner similar to VSG-treated control mice (Figure J-L). Thus, VSG normalized body weight and fat mass in LCK^TetTox^ mice and promoted similar improvements in indices of glucose homeostasis in LCK^TetTox^ and control mice.

These findings suggest that, despite the important roles for these cells in the physiologic restraint of feeding and body weight, neurons other than NTS^LCK^ cells must mediate the response to VSG. To determine whether additional NTS neurons might contribute to the refeeding response in control animals and/or in mice subjected to VSG, we quantified the FOS response to refeeding in control or VSG animals expressing reporter fluorophores in NTS^LCK^ cells (Supplemental Figure 4). This analysis revealed that NTS^LCK^ cells represent approximately 20% of feeding-activated NTS neurons in control mice, and approximately 35% of such neurons following VSG. Thus, in addition to neurons in the AP, hypothalamus, and elsewhere, it remains possible that non-NTS^LCK^ feeding-stimulated neurons within the NTS contribute to the response to VSG (as well as to other aspects of food intake suppression and body weight control).

## Discussion

Our findings reveal that distinct groups of food intake-inhibiting NTS neurons act additively to suppress food intake under physiologic feeding as well as during artificial activation. We also show that the enhanced suppression of food intake mediated by the activation of multiple populations of individually non-aversive NTS neurons does not promote aversive effects irrespective of the strength of food intake suppression, consistent with the notion that the pathways that promote the non-aversive suppression of food intake are distinct from those that promote aversion. Finally, Lepr^NTS^, Calcr^NTS^, and Cck^NTS^ neurons individually and coordinately restrain body weight gain during HFD feeding but are dispensable for weight loss in response to VSG, suggesting roles for additional neuron types in feeding and the response to VSG.

*Calcr* and *Lepr* expression do not colocalize in the same NTS neurons and unbiased clustering methods reveal that Calcr^NTS^ and Lepr^NTS^ neurons map to distinct populations of NTS neurons in mice (Calcr^NTS^ neurons map to GLU11, while Lepr^NTS^ neurons map mainly to GLU13 and GLU2; GLU13 controls feeding) (5, 6, 17, 20). Each of these populations strongly suppresses food intake but fails to promote a CTA when activated, suggesting that these NTS cells activate pathways that blunt feeding without engaging distinct aversive pathways (4, 6, 20). Our finding that the combined activation of these two cell types completely abrogates feeding over the first 24 hours of treatment, but fails to promote a CTA, further reinforces the notion that brainstem pathways need not provoke aversive responses to suppress food intake.

The magnitude of the anorexia that accompanies the combined activation of Calcr^NTS^ and Lepr^NTS^ cells relative to the response to the activation of a single population also reveals that distinct populations of food intake-suppressing NTS neurons can act additively to suppress feeding more effectively. Similarly, while the silencing of either of these populations can increase physiologic food intake and body weight in HFD-fed animals, silencing multiple populations provokes greater feeding and body weight gain. Hence, the various sets of food intake-suppressing NTS neurons must mediate additive effects to suppress physiologic feeding, at least under HFD-fed conditions.

The finding that silencing Calcr^NTS^, Lepr^NTS^, and Cck^NTS^ cells increases food intake and body weight in HFD-fed animals, but barely (if at all) alters these parameters in chow-fed animals indicates the importance of these cells for the restraint of food intake specifically during the consumption of palatable calorically-dense food. This might indicate that these NTS cell types play more prominent roles in the nutrient-(rather than volume/gut stretch) based suppression of feeding; alternatively, it might indicate that NTS neurons such as these play a stronger role in limiting hedonic overeating than in the restraint of physiologic, need-based feeding. Disentangling these possibilities will require a substantial amount of future work. It is also possible that NTSCck neurons play additional roles (other than the suppression of food intake) in the control of energy balance.

The finding that Calcr^NTS^, Lepr^NTS^, and/or Cck^NTS^ cell activation blocks feeding and promotes weight loss and that these cells play prominent roles in the suppression of food intake and body weight in HFD-fed animals suggested that they might also play crucial roles in the weight-loss response to VSG. Indeed, VSG increased the activation of these cells in response to an intragastric food bolus. Silencing these NTS neurons failed to blunt the VSG-mediated decrease in food intake and body weight, however. Hence, the action of these NTS neuronal populations is not required for the response to VSG. This indicates that other neural systems must mediate most of the weight-loss response to VSG. While the neurons that mediate this response to VSG might lie outside of the NTS (e.g., in the AP and/or hypothalamus), the finding that feeding and VSG activate many NTS neurons other than Calcr^NTS^, Lepr^NTS^, and/or Cck^NTS^ cells suggests that additional NTS populations might also contribute to the VSG response.

Although many sets of NTS neurons have been implicated in the suppression of feeding, only recently have single cell RNA sequencing methods been employed to define populations of NTS neurons in an unbiased manner (2, 5, 21, 22). Unlike *Calcr, Lepr* marks multiple populations of NTS neurons, including one population (GLU2) that controls respiratory function in response to leptin/energy balance (5, 22, 23). Also, *Cck* expression distributes across many informatically-defined NTS populations (including some Lepr^NTS^ cells), and it remains unclear which *Cck*-expressing NTS snRNA-seq population(s) might mediate the aversive suppression of food intake. Furthermore, *Calcr* and *Lepr* do not mark all neurons within their bioinformatically-defined populations (GLU11 and GLU13, respectively); the other neurons within these populations presumably mediate similar functions but are not altered by *Calcr*-or *Lepr*-directed manipulations, respectively.

Hence several sets of non-Calcr^NTS^, Lepr^NTS^, Cck^NTS^ cells might contribute to the ongoing control of food intake and the response to VSG in mice where the Calcr^NTS^, Lepr^NTS^, and/or Cck^NTS^ cells are silenced. These could include: the remaining unperturbed neurons from GLU11 and/or GLU13; the presumptive aversive NTS neuron population marked by *Cck* expression; other populations of NTS neurons that mediate the aversive and/or non-aversive suppression of food intake; and/or neuronal populations that lie outside of the NTS (including in the AP and/or hypothalamus). Going forward, it will be important to identify and study all the NTS cells that respond to food intake and VSG, as well as to identify neurons in the AP and elsewhere that may contribute to energy balance and the response to VSG.

## Materials and Methods

### Animals

Mice were bred in our colony in the Unit for Laboratory Animal Medicine at the University of Michigan; these mice and the procedures performed were approved by the University of Michigan Committee on the Use and Care of Animals and in accordance with Association for the Assessment and Approval of Laboratory Animal Care and National Institutes of Health guidelines. Mice were provided with food and water *ad libitum* (except as noted below) in temperature-controlled rooms on a 12-hour light-dark cycle. For all studies, animals were processed in the order of their ear tag number, which was randomly assigned at the time of tailing (before genotyping). ARRIVE guidelines were followed; animals were group-housed except for feeding and CTA studies.

We purchased male and female C57BL/6 mice for experiments and breeding from Jackson Laboratories. *Lepr*^*cre*^, *Calcr*^*cre*^, and *Cck*^*cre*^ mice have been described previously (24, 25) and were propagated by intercrossing homozygous mice of the same genotype. *Cck*^*cre*^ mice (Jax stock No.: 012706) for breeding were purchased from Jackson Labs (Bar Harbor, ME).

### Stereotaxic Injections

AAV^Flex-hM3Dq^, AAV^Flex-hM4Di^ and AAV^Flex-TetTox-GFP^ were prepared by the University of Michigan Viral Vector Core. The AAVs used in the manuscript were all serotype AAV8. For injection, following the induction of isoflurane anesthesia and placement in a stereotaxic frame, the skulls of adult mice were exposed. After the reference was determined, a guide cannula with a pipette injector was lowered into the injection coordinates (NTS: A/P, −0.2; M/L, ±0.2; D/V, −0.2 from the obex) and 100 nL of virus was injected for each site using a picospritzer at a rate of 5-30 nL/min with pulses. Five minutes following injection, to allow for adequate dispersal and absorption of the virus, the injector was removed from the animal; the incision site was closed and glued. The mice received prophylactic analgesics before and after surgery. The mice injected with AAV^Flex-hM3Dq^, AAV^Flex-hM4Di^, AAV^Flex-TetTox-GFP^, or control viruses were allowed at least 1 week to recover from surgery before experimentation.

### Phenotypic studies

LepRb^TetTox^, Cck^TetTox^, LC^TetTox^, and LCK^TetTox^ mice and their controls were monitored from the time of surgery for chow feeding. For stimulation studies, DREADD-expressing mice and their controls that were at least three weeks post-surgery were treated with saline or drug (CNO, 4936, Tocris) at the onset of dark cycle, and food intake was monitored over four hours. For chronic food intake and body weight changes in DREADD-expressing animals, mice were given saline for two to three days prior to injecting saline or drugs twice per day (approximately 5:30 PM and 8:00 AM) for 2 or 3 days, followed by saline injections for another one or three days to assess recovery from the treatment.

### Perfusion and immunohistochemistry

Mice were anesthetized with a lethal dose of pentobarbital and transcardially perfused with phosphate-buffered saline (PBS) followed by 10% buffered formalin. Brains were removed, placed in 10% buffered formalin overnight, and dehydrated in 30% sucrose for 1 week. With use of a freezing microtome (Leica, Buffalo Grove, IL), brains were cut into 30 μm sections. Immunofluorescent staining was performed using primary antibodies (FOS, #2250, Cell Signaling Technology, 1:1000; GFP, GFP1020, Aves Laboratories, 1:1000; dsRed, 632496, Takara, 1:1000), antibodies were reacted with species-specific Alexa Fluor-488, −568 or −647 conjugated secondary antibodies (Invitrogen, Thermo Fisher, 1:200). Images were collected on an Olympus (Center Valley, PA) BX53F microscope. Images were pseudocolored using Photoshop software (Adobe) or Image J (NIH).

### Conditioned taste aversion (CTA)

Two types of CTA assay were conducted, as described previously (6): Saccharin CTA: Mice that were 8 weeks of age or older were individually housed in standard cages with low wire tops and free access to food and the lixits were removed. Mice were habituated to two water-containing bottles for 3-5 days until they learned to concentrate their daily water consumption into these two water bottles. On the conditioning day, the mice received only two saccharin (0.15%, 240931, Sigma) bottles. Following the 60 min exposure to saccharin, mice were injected intraperitoneally with the desired stimulus (vehicle control (0.9% NaCl), lithium chloride (0.3 M, 203637, Sigma)) in a volume equivalent to 1% of each animal’s body weight (10 mL/g), or CNO (1 mg/kg, 4936, Tocris Bioscience) Access to the two saccharin bottles continued for an additional 1 h, followed by the return of normal water bottles. Two days later, each mouse received access to two water bottles (one containing 0.15% saccharine, the other containing water), and the amount of fluid ingested from each water bottle was be measured.

HFD CTA: following an overnight fast, mice that were 8 weeks of age or older were provided with one hour access of HFD (D012492, Research Diets), followed with the relevant stimuli, followed by an extra one hour of HFD access before returning of normal chow. On the post-conditioning day, fasted mice received access to both HFD and chow and the consumption of each was measured.

### Mouse Vertical Sleeve Gastrectomy

Mice had *ad libitum* access to water and 60% lard-based HFD (Research Diet, New Brunswick, NJ; Cat. No. D12492) for 6-8 weeks prior to undergoing Sham or VSG surgery. Under isoflurane/O_2_ mixture anesthesia, all mice received a midline incision in the ventral abdominal wall and the stomach was exposed. For VSG, approximately 80% of the stomach was transected along the greater curvature using an Echelon Flex Powered Vascular 35mm Stapler Model (PVE35A; Ethicon endo-surgery) creating a gastric sleeve. The Sham surgery was performed by the application of gentle pressure on the stomach with blunt forceps for 15 seconds. All mice received one dose of Buprinex (0.1 mg/kg) and Meloxicam (0.5 mg/kg) immediately after surgery. Post-operatively, all mice received 1 ml warm saline subcutaneously on the first postoperative day and analgesic treatment with meloxicam (0.5 mg/kg) for 3 consecutive days. Animals were placed on DietGel Boost (ClearH_2_O; Postland ME) for 2 days before and 3 days after surgery before pre-operative solid diet (60% HFD) was returned on day 4. Body weight and general health were observed daily for the first 10 days post-surgery. Food intake and body weight were measured weekly for the duration of the study. Body composition was measured using nuclear magnetic resonance (EchoMRI^™^-900, EchMRI LLC, Houston, USA) before surgery and periodically throughout the study.

### Glucose/Insulin/Acetaminophen assay

During glucose tolerance tests, all mice were fasted for 4-5 hours prior to oral administration of dextrose (2 g/kg body weight). Tail vein blood glucose levels were measured using Accu-Chek glucometers (Accu-Chek Aviva Plus, Roche Diagnostics) or Biosen C-line glucose analyzer (EKF diagnostic) at 0, 15, 30, 45, 60, 90, and 120 minutes post glucose administration. Gastric emptying rate was assessed by an oral gavage of glucose (2 g/kg) with acetaminophen (100mg/kg) in 4-5 hour fasted mice. Blood was collected from the tail vein at baseline and 10 min after gavage in EDTA-coated microtubes. Plasma acetaminophen levels were used to assess the rate of gastric emptying and were measured using spectrophotometry assay (Sekisui Diagnostics). Plasma insulin levels were determined using ELISA colorimetric insulin assay kit (Crystal Chem).

### Metabolic Chamber

A subset of mice was placed into an automated system equipped with metabolic chambers (TSE Systems International Group, Chesterfield, MO) to measure indices of energy homeostasis 5–7 wk after surgery. Data were recorded continuously over the course of 7 days, and the last 4 days were used in the analysis.

### Statistics

Statistical analyses of the data were performed with Prism software (version 8). Two-way ANOVA, paired or unpaired t-tests were used as indicated in the text and figure legends. In detail, multiple groups with one variable comparison were performed using one-way ANOVA post hoc Tukey. Multiple groups with two variables comparison were performed using two-way ANOVA post hoc Sidak. Multiple group comparisons over time were performed using repeated measures ANOVA post hoc Tukey. All data are presented as Mean ± SEM. Two groups direct comparisons were performed using t-test. Two groups multiple comparisons were also performed using Multiple t-test with Bonferroni correction. Data were considered statistically significant when P<0.05.

## Acknowledgments

We thank Paula Goforth, PhD, David Olson, MD, PhD, and Randy Seeley, PhD, as well as members of the Myers and Sandoval labs for helpful discussions. Special thanks to Stace Kernodle and Kelli Rule for superb technical support.

## Author contributions

Conceptualization, WQ, CH, MGM, and DS; Investigation, WQ, CH, YW, and JW; Writing, WWC, CH, YW, JW, MGM, and DS; Funding Acquisition, MGM and DS.

## Data and materials availability

All data will be freely available upon publication. Mouse models and reagents will be made available to academic laboratories.

## Supplemental Figures

**Supplemental Figure 1:**
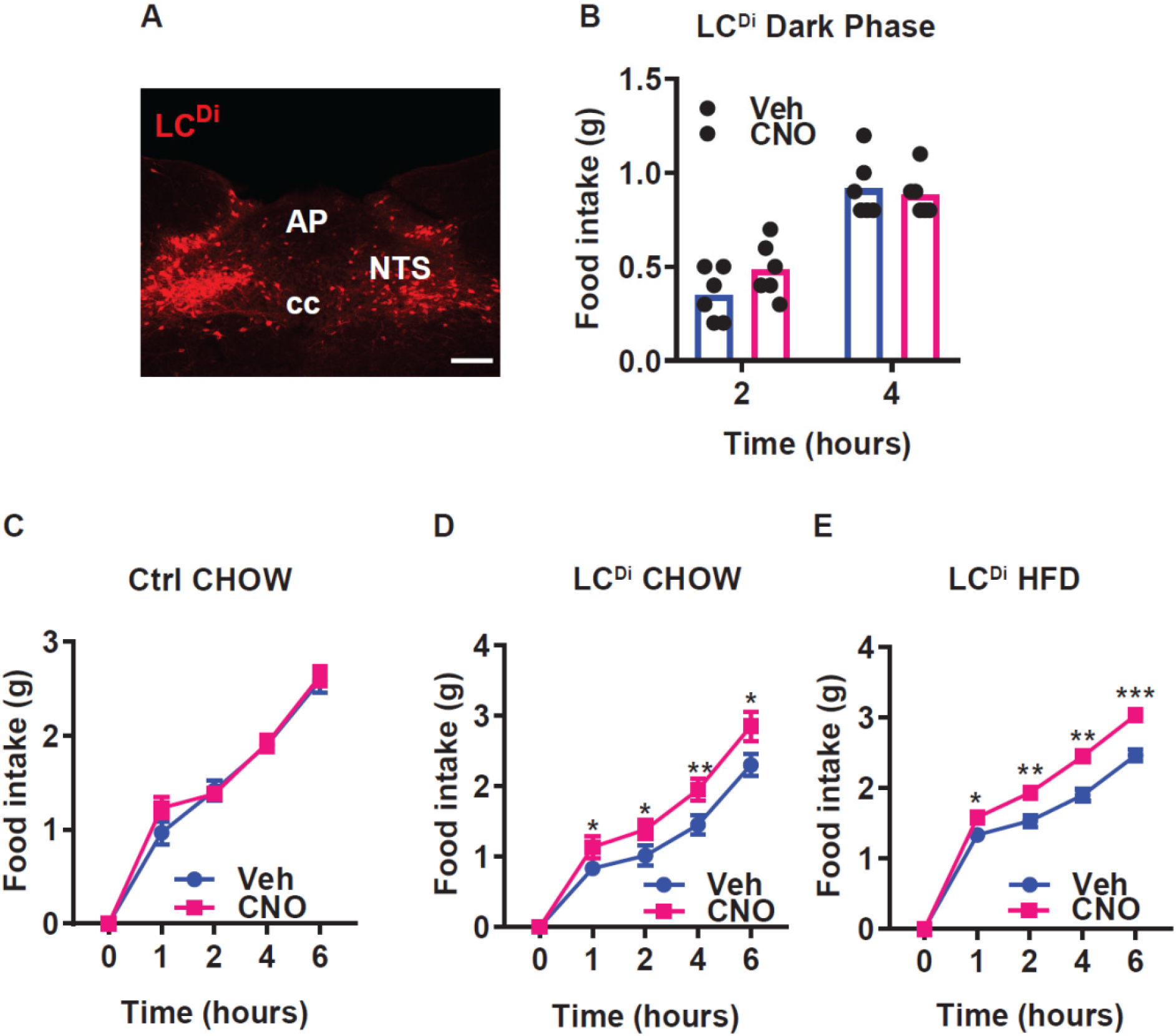
Inhibiting LC^NTS^ neurons increases food intake. **(A)** Representative NTS image showing dsRed-IR (red, mCherry) in the NTS of *Lepr*^*Cre*^*;Calcr*^*Cre*^ mice injected with mCherry-expressing AAV^Flex-hM4Di^ (LC^Di^ mice). NTS: the nucleus of the solitary tract, AP: area postrema, cc: central canal; scale bar equals 150 μm. (**B**) Food intake in LC^Di^ mice during treatment with vehicle (Veh) or CNO injection (IP, 1 mg/kg) at the onset of dark cycle, n=6 in each group. **(C)** Food intake during the first 6 hours of refeeding in the light cycle following an overnight fast for control (**C**, n=6 per condition) or LC^Di^ (**D**, n=6 per condition) animals fed with chow and for LC^Di^ animals fed with HFD (**E**, n=6 per group) during treatment with CNO (IP, 1 mg/kg) or Veh. Shown is mean +/-SEM. Two-way ANOVA, sidak’s multiple comparisons test was used, *p<0.05, **p<0.01, ***p<0.001 vs Veh.

**Supplemental Figure 2:**
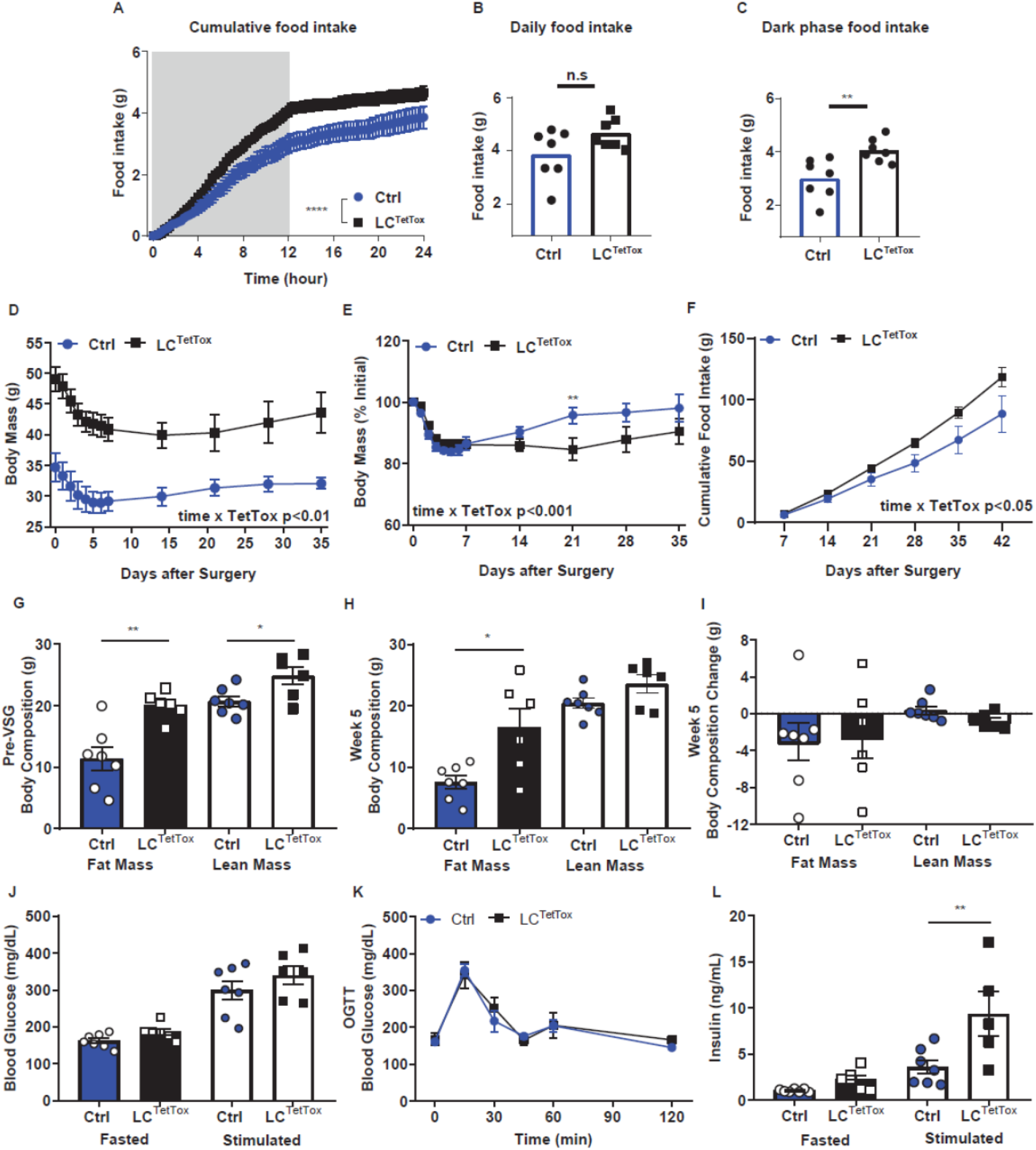
Silencing NTS^LC^ neurons increases food intake and body weight but fails to abrogate the weight loss effects of VSG. (**A-C**) Cumulative food intake (**A**), daily food intake (**B**) and dark phase food intake (**C**) were continuously monitored over 24 hours in a TSE system during the 7th week after surgery with control (Ctrl, n=7) and LC^TetTox^ (n=7) animals. (**D-L**) Response to VSG for Ctrl and LC^TetTox^ mice. Body mass (absolute (**D**), and %initial (**E**)), cumulative food intake (**F**), body composition before VSG (**G**), 5 weeks after VSG (**H**) and change in body composition (**I**) are shown for Ctrl and LC^TetTox^ mice following VSG. (**J-L**) Response to an oral glucose load (**J**, 2 g/kg, gavage), along with fasting and stimulated (10 minutes after gavage) glucose (**K**) and insulin (**L**) concentrations are shown for Ctrl and LC^TetTox^ mice following VSG. Shown is mean +/-SEM. For A-C, two-way ANOVA, sidak’s multiple comparisons test was used. For **D-L**, data was analyzed using RM-ANOVA or Student’s T-test; **p<0.01, ****p<0.0001 vs Ctrl. Mice were a mixture of males and females; Ctrl, n=7; LC^TetTox^, n=6.

**Supplemental Figure 3:**
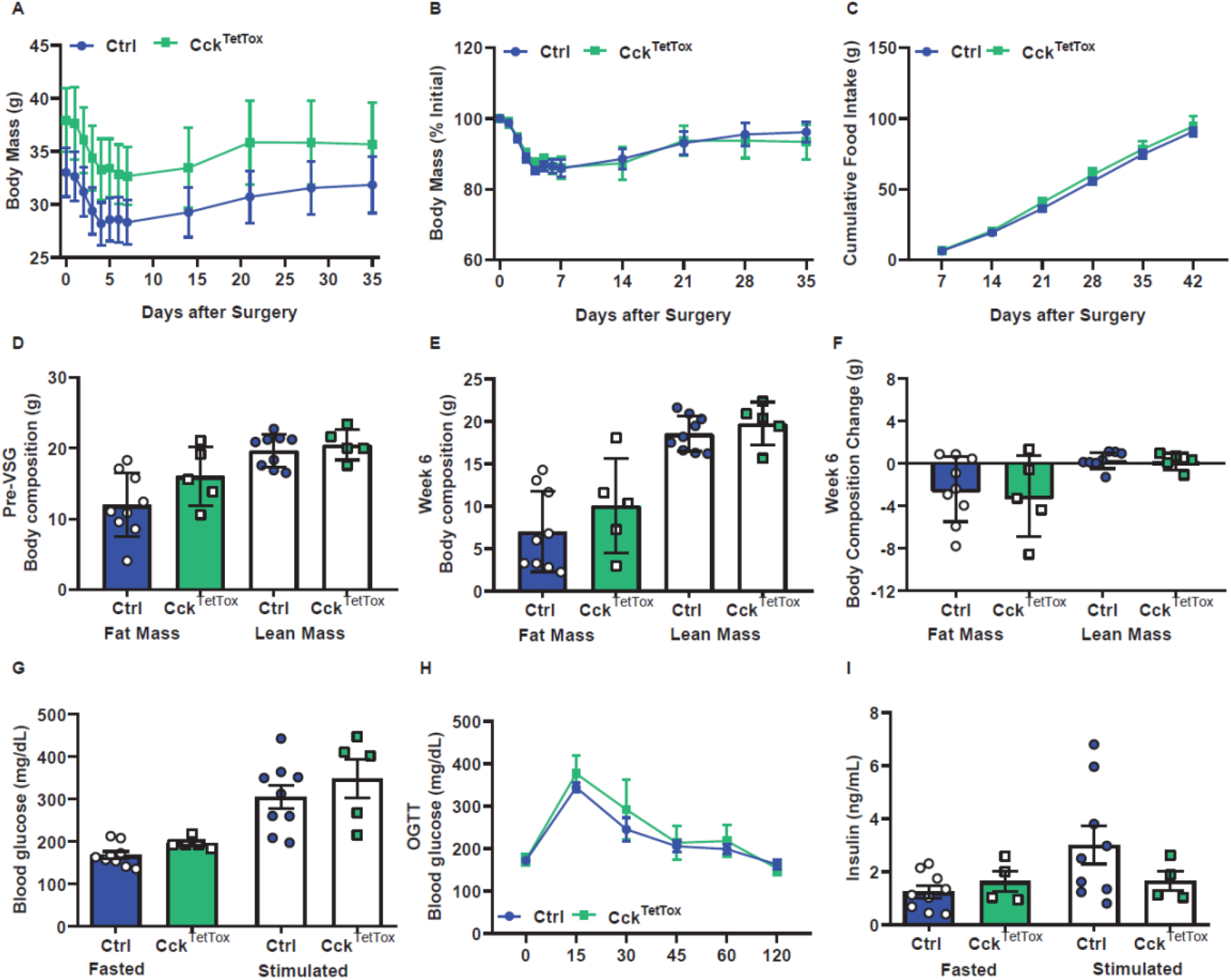
Response of Cck^TetTox^ mice to VSG. Body mass (absolute (**A**), and %initial (**B**)), cumulative food intake (**C**), body composition before VSG (**D**), 5 weeks after VSG (**E**) and change in body composition (**F**) are shown for Ctrl and Cck^TetTox^ mice following VSG. (**G-I**) Response to an oral glucose load (**G**, 2 g/kg, gavage), along with fasting and stimulated (time) glucose (**H**) and insulin (**I**) concentrations are shown for Ctrl and Cck^TetTox^ mice following VSG. Shown is mean +/-SEM. Data was analyzed using RM-ANOVA or Student’s T-test. Mice were a mixture of males and females (Ctrl, n=9; Cck^TetTox^, n=4-5).

**Supplemental Figure 4:**
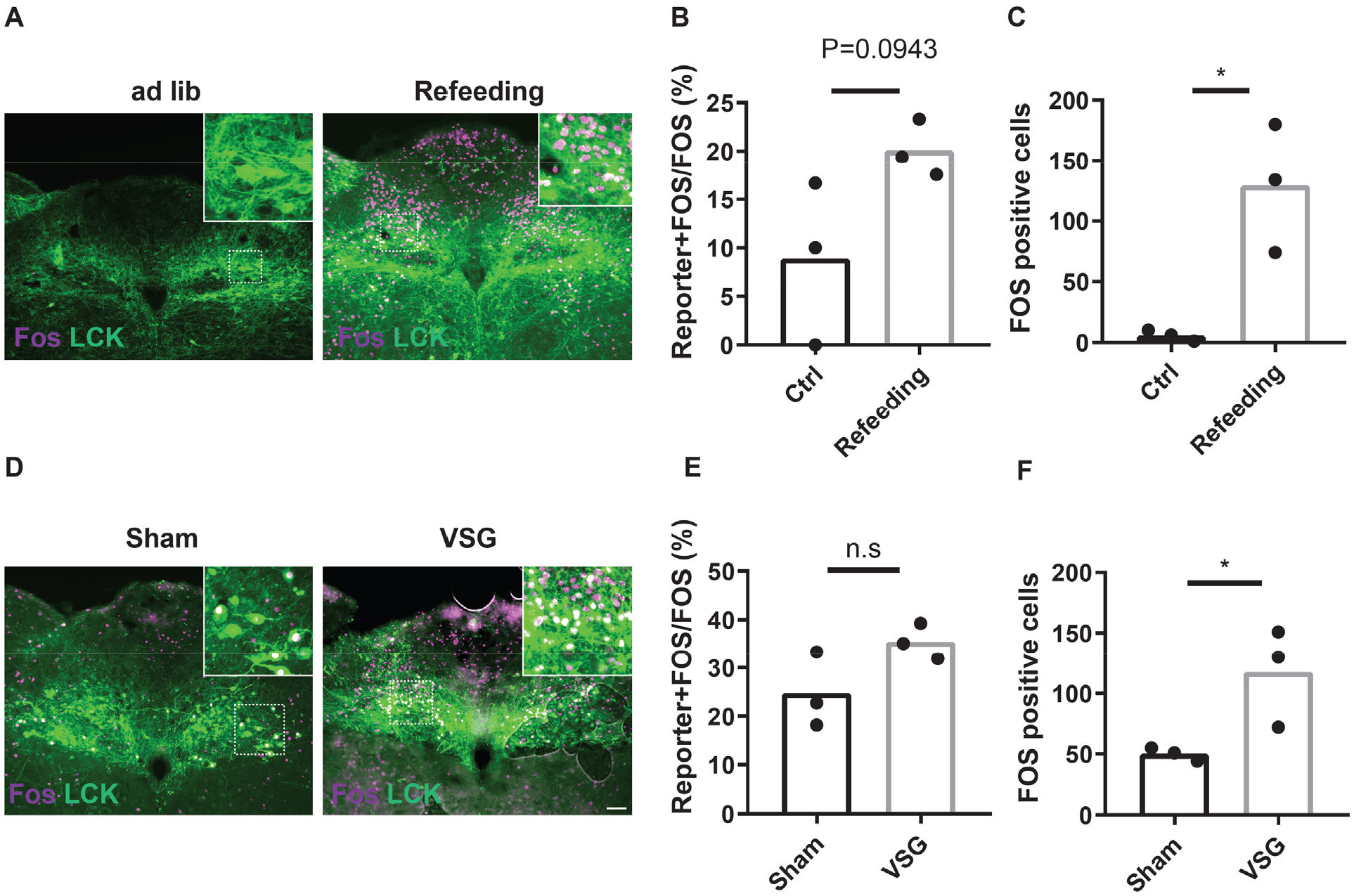
Activation of NTS^LCK^ and other NTS neurons in response to refeeding and VSG. *Lepr*^*Cre*^*;Calcr*^*Cre*^*;Cck*^*Cre*^ mice were injected with cre-dependent AAV reporters (LCK^Reporter^ mice). LCK^Reporter^ mice were treated as described below and stained for the appropriate reporter (LCK, green) and FOS (magenta). **A-C**: LCK^Reporter^ mice were fed *ab libitum* (Ctrl) or overnight fasted (16 hours) and refed (Refeeding) prior to perfusion. (**A**) Representative images of the NTS. (**B-C**) Quantification of Reporter+FOS colocalized neurons as a percentage of total FOS neurons (**B**) and total FOS neurons (**C**) per section. (**D-F**) LCK^Reporter^ mice were subjected to sham surgery (Sham) or VSG. After recovery, they were fasted for 4 hours during the light cycle and gavaged with a glucose load prior to perfusion. (**D**) Shows representative images of the NTS. (**E-F**) Quantification of Reporter+FOS colocalized neurons as a percentage of total FOS neurons (**B**) and total FOS neurons (**C**) per section. Shown is mean +/-SEM; n=3 mice per group. Student’s unpaired t-test was performed. n.s= not significantly different; * p<0.05 vs Ctrl or Sham. Scale bar equals 150 μm.

